# STAYGREEN-mediated chlorophyll a catabolism is critical for photosystem stability upon heat stress in ryegrass

**DOI:** 10.1101/2021.09.10.459836

**Authors:** Jing Zhang, Hui Li, Xinru Huang, Jing Xing, Jiaming Yao, Jiafu Jiang, Puchang Wang, Bin Xu

## Abstract

Chlorophyll (Chl) loss is one of the most visible symptoms of heat-induced leaf senescence, especially for cool-season grass species. Suppression of the Chl a Me-dechelatase gene, *SGR* (also named as *nye1*), blocked the degradation of Chl a and resulted in the ‘stay-green’ trait during leaf senescence. However, effect of Chl a catabolism on plant tolerance to long-term moderate heat stress (35-40°C) remains unclear. In this study, we suppressed the expression of Chl a catabolic gene, *LpSGR*, in both constitutive and inducible manners in perennial ryegrass. Constitutive suppression of *LpSGR* aggravated heat stress-induced chloroplast structure and photosystem damages, disrupted energy utilization/dissipation during photosynthesis, activated ROS generation with weakened ROS-scavenging enzyme activities. Transcriptome comparison among wildtype (WT) and transgenic RNAi plants under either the optimum or high temperature conditions also emphasized the effect of Chl a catabolism on expression of genes encoding photosynthesis system, ROS-generation and scavenging system, and heat shock transcription factors. Furthermore, making use of a modified ethanol-inducible system, we generated stable transgenic perennial ryegrass to suppress *LpSGR* in an inducible manner. Without ethanol induction, these transgenic lines exhibited the same growth and heat tolerance traits to WT, while under the induction of ethanol spray, the transgenic lines also showed compromised heat tolerance. Taken together, our data suggest that Chl a catabolism is critical for energy dissipation and electron transfer in photosynthesis, ROS-balancing and chloroplast membrane system stability upon long-term moderate heat stress.

## Introduction

Chlorophyll (Chl) molecules, mainly Chl a & Chl b, are key pigments in absorbing light quantum and transferring the energy to the photosynthesis reaction center (Tanaka and Tanaka, 2006; Morita et al., 2009). It is more efficient for Chl a to absorb short-wavelength light than Chl b, and the energy of photon is higher when the light wavelength is shorter. Upon the absorption of higher energy photon, Chl transits to a higher-energy, or excited, state (Chl*) (Taiz and Zeiger, 2002). The energy stored in Chl* is rapidly dissipated by excitation transfer or photochemistry. If the Chl* is not rapidly quenched, it can react with molecular oxygen to form an excited state of oxygen known as singlet oxygen (^1^O_2_) that reacted and further produces reactive oxygen species (ROS, e.g., O_2_^−^ and H_2_O_2_) (Wilson et al., 2006). The overproduced ROS goes on to react with and damage many cellular components, especially lipids. In turn, one way that plants use to suppress this phototoxicity efficiently is through Chl catabolism (Liu and Guo, 2013).

Under the natural habitat, light and temperature are determinant factors for plants to survive, being in a state of flourish, deciduous, dormant or dying. Long term moderately high temperature (35-40°C) reduces photosynthesis rate. Photosystem Ⅱ (PSⅡ) is the most heat sensitive system that it is inhibited at lower temperature than that needed to damage it (Sharkey, 2005). Heat stress triggers ROS (e.g., H_2_O_2_) production in cells mainly due to the energy dissipation during electron transfer and Chl* quenching in photosynthesis as mentioned above. ROS damages lipid and other cellular systems in one hand but also serves as signal molecules to activate heat stress proteins (e.g., HSF) and heat-responsive genes’ expression (Li et al., 2021), among which Chl catabolic genes were the representative ones (Jespersen et al., 2016; Yu et al., 2021). And it is known that dismantling Chl from PSⅡ can cause the system being photosensitive followed by rapid degradation of PSⅡ in a whole (Hirashima et al., 2009; Li et al., 2017). In short, Chl are key molecules in the regulation of PSⅡ’s efficiency and integrity as well as ROS’ generation upon heat stress. Yet, how Chl metabolism may affect plant heat tolerance is not well understood.

Upon long-term moderate heat stress (e.g., 35-40°C), cool season grasses visually responded in a way with reduced leaf Chl content (i.e., leaf yellowing), and this reduced Chl content was mainly due to accelerated Chl degradation but not inhibited Chl biosynthesis (Jespersen et al., 2016). Despite of the common conclusion that accelerated Chl catabolism is a beneficial process to suppress this heat-induced phototoxicity, we found that suppressing a Chl b reductase (*NOL*) gene’s expression to inhibit Chl b degradation improved, instead of undermined, plant heat tolerance in a cool-season grass, perennial ryegrass (*Lolium perenne*) (Yu et al., 2021). This finding leads us to hypothesize that not only the whole process of Chl catabolism, but specific steps of the process mattered in the regulation of plant heat tolerance. Chl a was converted to pheophytin a catalyzed by (Mg)-dechelatase encoded by *STAYGREEN* [*SGR*, or *NON-YELLOING 1* (*NYE1*)] (Shimoda et al., 2016). As mentioned above, Chl a absorbed more higher energy light quantum than Chl b that can lead to the generation of Chl*. It is possible that some plants cope with moderate long-term heat stress by preferentially down-regulating Chl a content to minimize Chl* and thereof ROS burst.

Perennial ryegrass is a cool-season perennial grass species widely cultivated in temperate regions throughout the world for its turf and forage purposes (Altpeter et al., 2000; Chastain et al., 2015). It is considered as a heat-sensitive species with early leaf-yellowing phenotype when suffered from heat stress. For this turf and forage plant species, it is highly desirable to retain the stay-green trait without compromised heat tolerance. Previously, stay-green *Lolium* species have been obtained through introgression of the *sid* locus containing the null *sgr* gene in meadow fescue into a *Lolium*/*Festuca* hybrid (Armstead et al., 2006; Thomas et al., 1997; Thomas et al., 1999). And transgenic ryegrass with suppressed *SGR* also retained the stay-green phenotype (Xu et al., 2019). To obtain stay-green and heat tolerance perennial ryegrass and to understand how Chl a catabolism may involve in plant heat tolerance, we suppressed the expression of Chl a catabolic gene, *SGR*, in both constitutive and inducible manners in perennial ryegrass.

## Results

### Knockdown of *LpSGR* lead to compromised plant heat tolerance and reduced tiller number in perennial ryegrass

Besides the typical stay-green trait as reported before (Xu et al., 2019), constitutive suppressing the expression of *LpSGR* lead to reduced tiller number as well longer but narrower leaf width in perennial ryegrass (table 1). For example, after one month of growth, the single tiller of WT proliferated into 11.0 tillers, while, two RNAi:*LpSGR* transgenic lines (*SGRi*), *SGRi*-1 and *SGRi*-6, only had 3.8 and 4.8 tillers, respectively, and therefore significantly lower aboveground biomass yield than those of WT (table 1).

**Table 1.**
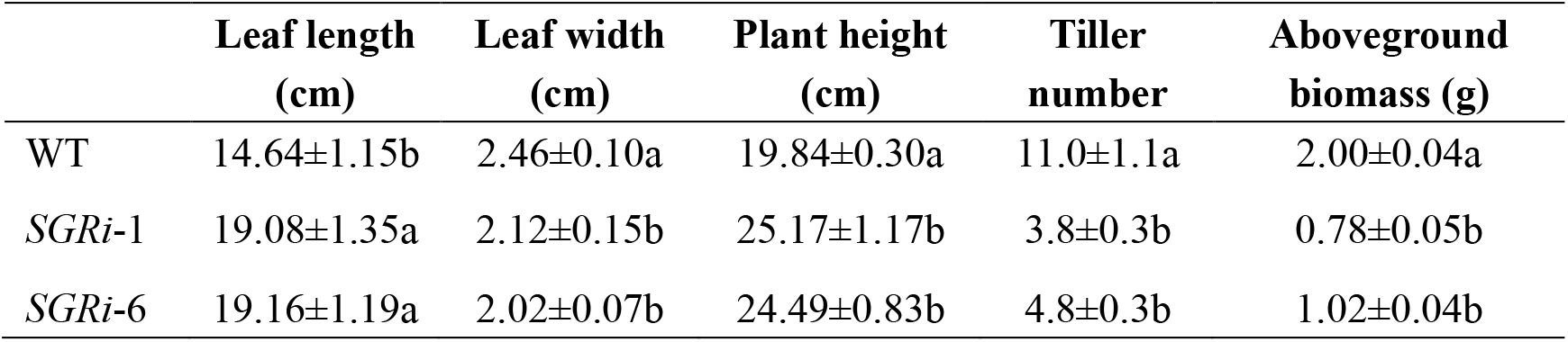
Effects of *LpSGR* knockdown on the aboveground growth of perennial ryegrass.

When exposed to prolonged heat stress for 21 days, the growth of *SGRi* lines were more severely affected than WT plants (Fig. 1A). Under the optimum growth condition, there was no significant difference between *SGRi* and WT plants for their Chl b content, net photosynthesis rate (Pn), electrolyte leakage (EL), and leaf relative water content (RWC) (Fig 1B-G). When under heat stress conditions, *SGRi* lines had significantly lower Chl a and b contents, Pn, and RWC, but higher Chl a/b ratio and EL than WT plants (Fig 1B-G).

**Figure 1.**
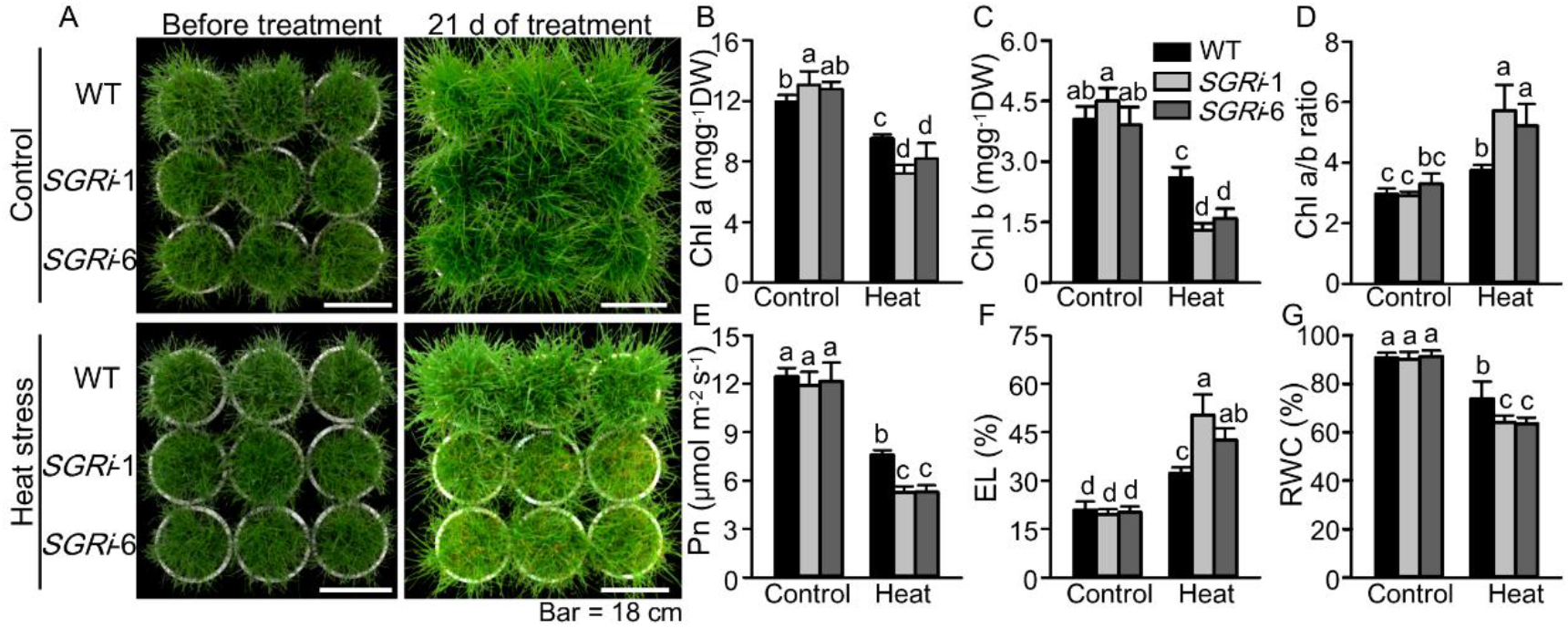
Phenotypic and physiological analysis of *SGRi* lines and WT plants under non-stress and heat stress conditions. (A) Phenotypes of WT and *SGRi* lines before and after 21 days of heat and non-stress treatment. The scale bars equal to 18 cm. (B-G) Chl a content (B), Chl b content (C), Chl a/b ratio (D), net photosynthesis rate (Pn) (E), electrolyte leakage (EL) (F), and leaf relative water content (RWC) (G). Data are means ± SE. Different letters above bars in each group represent significant difference at p ≤ 0.05.

### Knockdown of *LpSGR* altered chloroplast ultrastructure and LHC protein

The chloroplast ultrastructure of *SGRi* lines was different from WT plants when grown under non-stress or heat stress conditions (Fig. 2). Under the optimum growth temperature, *SGRi* lines had higher number of grana *per* chloroplast, smaller grana size, and more tightly stacked grana thylakoids than those of WT (Fig. 2A-C). Under heat stress, the number and size of plastoglobule (PG) was significantly increased by heat stress treatment in both *SGRi* lines and WT plants, while the chloroplast and thylakoid membranes were more severely degraded, the grana stacks were thicker, and intergrana thylakoids were fewer in *SGRi* lines than those of WT plants (Fig. 2D-F).

**Figure 2.**
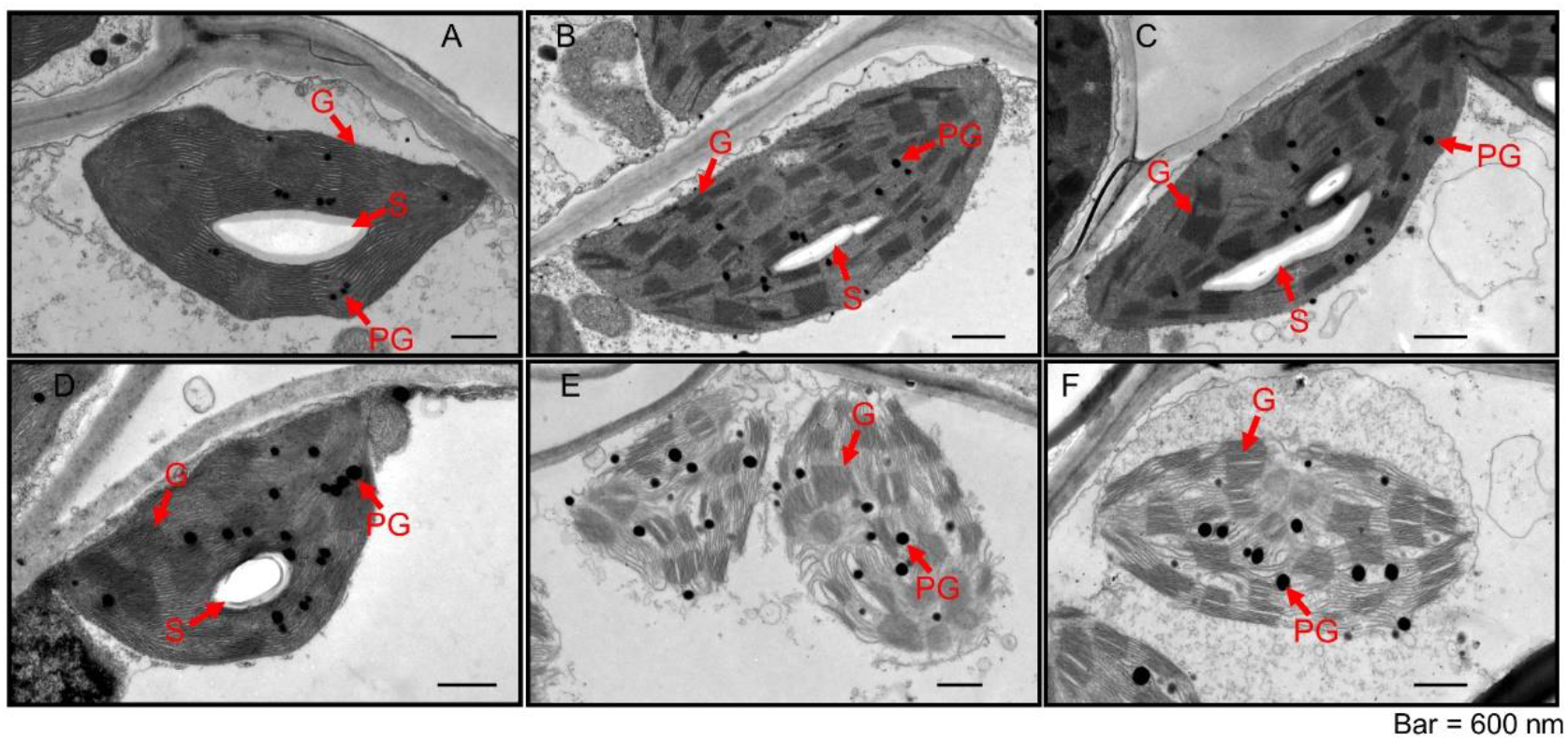
Transmission electron microscopy of chloroplast in *SGRi* lines and WT plants under non-stress and heat stress conditions. (A-C) Chloroplasts in the mesophyll cells of 3^rd^ leaves form the top in WT (A), *SGRi*-1 (B), and *SGRi*-6 (C) at 21 days of non-stress treatment. (D-F) Chloroplasts in the mesophyll cells of 3^rd^ leaves form the top in WT (D), *SGRi*-1 (E), and *SGRi*-6 (F) at 21 days of heat stress treatment. G, grana thylakoid; PG, plastoglobule; S, starch. At least 20 chloroplasts were observed in each plant. The scale bars equal to 600 nm.

Consistent to the chloroplast ultrastructure observation, immunoblotting analysis also showed that heat stress significantly decreased the abundance levels of photosynthesis proteins which are involved in light harvesting, electron transport, and carbon dioxide assimilation, such as Lhca3, Lhcb1/2/3/5, PsaA, PsbA (D1), PsbD (D2), and RbcL, and such protein abundance decreases were more severe in *SGRi* than in WT plants (Fig. 3A). The total protein content of leaves of *SGRi* lines were also significantly lower in leaves of *SGRi* lines than those in WT plants under heat stress condition (Fig. 3B).

**Figure 3.**
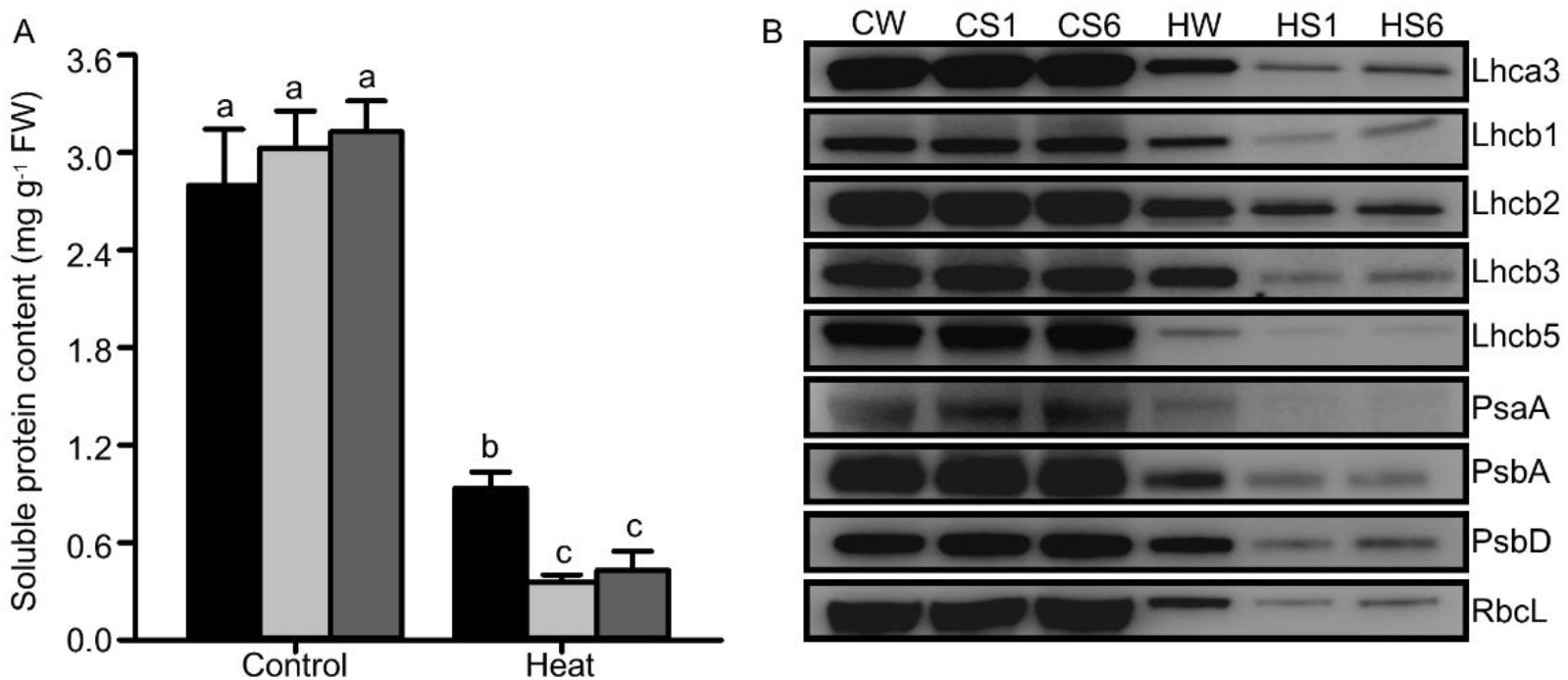
Immunoblot of photosynthesis proteins in *SGRi* and WT plants under non-stress and heat stress. (A) The protein content of leaves from WT and *SGRi* lines under non-stress control and heat stress conditions. (B) Immunoblot of PSⅠ antenna (Lhca3), PSⅠ core (PsaA), PSⅡ antenna (Lhcb1, Lhcb2, Lhcb3, and Lhcb5), PSⅡ core (PsbA and PsbD), and RbcL (Rubisco large subunit). This experiment was repeated three times with similar results. Data are means ± SE. Different letters above bars in each group represent significant difference at p ≤ 0.05.

### Knockdown of *LpSGR* affected photosynthetic electron transport rate and quantum yield of PSⅡ

Knockdown of *LpSGR* also affected light energy absorption and utilization for photochemical electron transport in a temperature-dependent manner. As shown in figure 4, under control condition, *SGRi* lines had similar values of the maximum photochemical efficiency of PSII (Fv/Fm), quantum yield of non-regulated energy dissipation [Y(NO)], and photochemical fluorescence quenching (qP) compared to those of WT plants. The quantum yield of PSⅡ [Y(Ⅱ)] and apparent photosynthetic electron transport rate (ETR) were significantly higher, but the quantum yield of regulated energy dissipation [Y(NPQ)] was significantly lower in *SGRi* lines than those of WT plants under non-stress conditions (Fig. 4). Yet, under heat stress, *SGRi* lines showed significantly lower Fv/Fm, Y(Ⅱ), qP, and ETR, but significantly increased Y(NPQ) than WT plants. In particular, *SGRi* lines showed significantly higher ETR and Y(Ⅱ), but lower Y(NPQ) under the optimum growth temperature, but the contrary was true for *SGRi* lines under heat stress, suggesting that knockdown of *LpSGR* affected photosynthetic electron transport rate and quantum yield of PSⅡ in a temperature-dependent manner.

**Figure 4.**
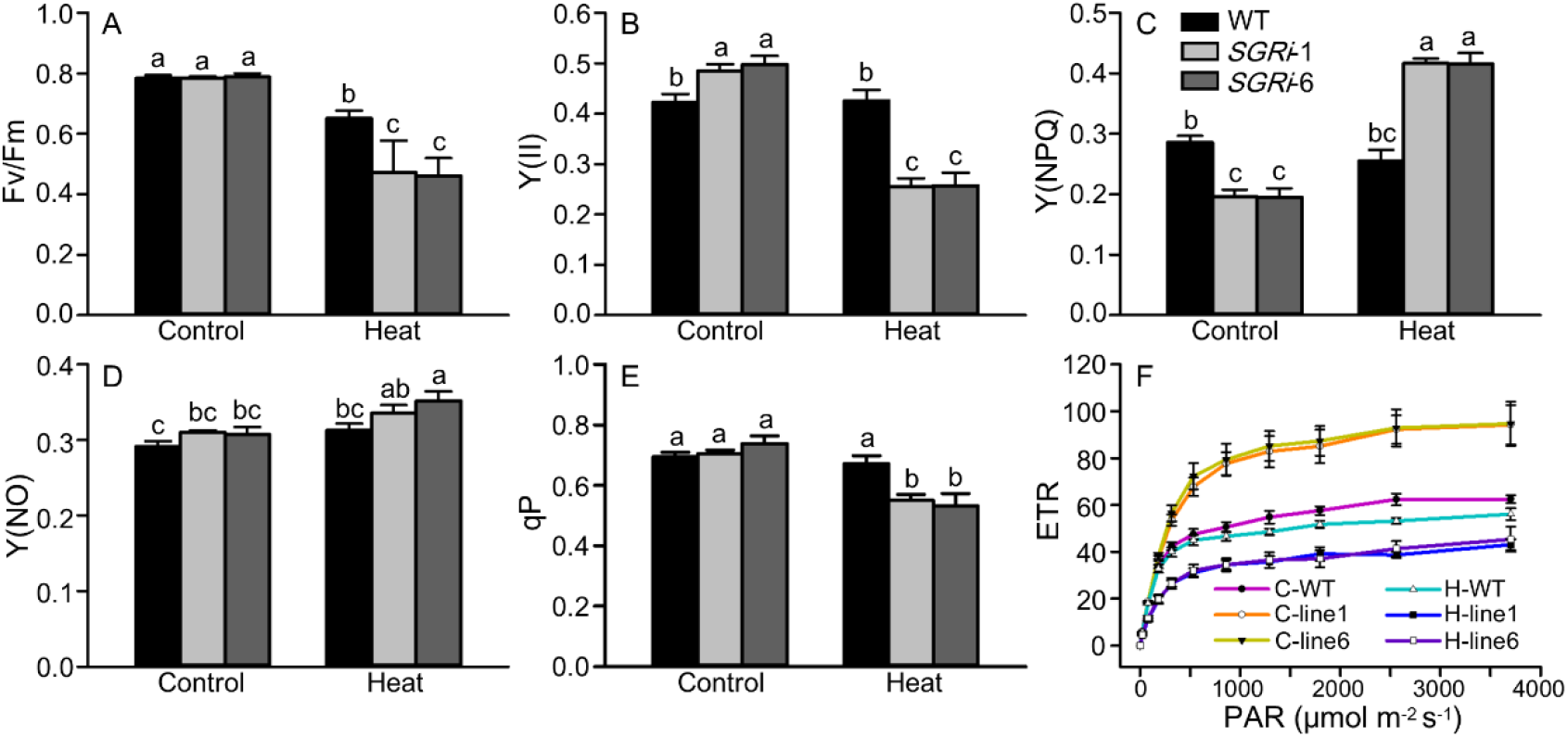
Chl fluorescence analysis in *SGRi* and WT plants under non-stress and heat stress conditions. (A-F) Maximum photochemical efficiency of PSII (Fv/Fm) (A), quantum yield of PSⅡ [Y( Ⅱ)] (B), quantum yield of regulated energy dissipation [Y(NPQ)] (C), quantum yield of non-regulated energy dissipation [Y(NO)] (D), photochemical fluorescence quenching (qP) (E), and apparent photosynthetic electron transport rate (ETR) (F), of WT and *SGRi* lines at 21 days of heat and non-stress treatment. Data are means ± SE. Different letters above bars in each group represent significant difference at p ≤ 0.05.

### RNA interference of *LpSGR* changed heat-induced reactive oxygen species (ROS) production, antioxidant enzyme activity and gene expression

Excessive energy generated from the electron transport chain under heat stress could lead to reactive oxygen species (ROS) accumulation, such as superoxide anions (O_2_^−^) and hydroxyl radicals. Therefore, we then compared the yield of ROS and activities of ROS-scavenging enzymes between *SGRi* lines and WT plants. As shown in figure 5, knockdown of *LpSGR* lead to significantly higher levels of O_2_^−^ and H_2_O_2_ under heat stress, but did not affect their yield when under the optimum temperature condition (Fig. 5A-D). Contrary to the accumulation of ROS, enzymatic activities of four kinds of ROS-scavenging enzymes (SOD, CAT, APX, and GPX) were lower in *SGRi* lines than WT plants under heat stress (Fig. 5F-H).

**Figure 5.**
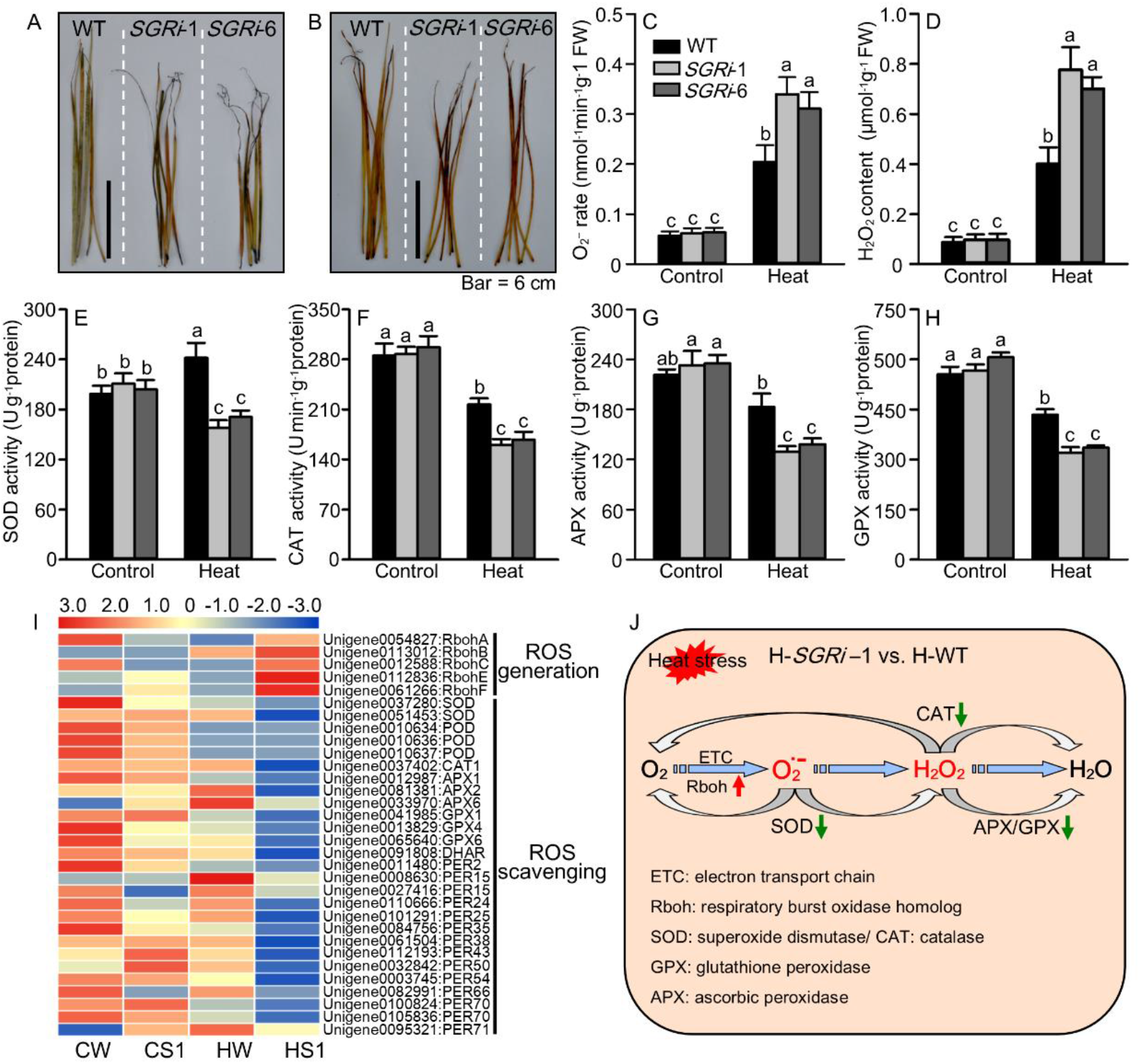
ROS production and scavenging enzymatic activities in *SGRi* and WT plants under non-stress and heat stress conditions. (A, B) Histochemical staining of O_2_^−^ (A) and H_2_O_2_ (B) in WT and *SGRi* leaves at 21 days of heat stress treatment. The scale bars equal to 6 cm. (C-D) O_2_^−^ production rate (C) and H_2_O_2_ content (D) of WT and *SGRi* leaves. (E-H) Activities of SOD (E), CAT (F), APX (G), and GPX (H) of WT and *SGRi* leaves. (I) Differential expressed genes (DEGs) annotated in ROS generation and scavenging pathways according to transcriptome comparison. The RPKM of each DEGs was used for drawing heatmap. Sample abbreviations: CW, WT under control condition; CS1, *SGRi*-1 under control condition; HW, WT under heat stress condition; HS1, *SGRi*-1 under heat stress condition. (J) The diagram for the DEGs annotated in ROS generation and scavenging pathways according to transcriptome of *SGRi*-1 compared to WT plants under heat stress condition (HS1 vs. HWT). Data are means ± SE. Different letters above bars in each group represent significant difference at p ≤ 0.05.

### Transcriptomic comparisons among *SGRi* and WT plants under optimum temperature or heat stress

Transcriptomes of *SGRi*-1 and WT plants were profiled from leaves at 21 days of heat stress treatment using Illumina sequencing. The contigs were assembled into 113,372 unigenes with N50 length of 1,514 bp, and an average unigene size of 895 bp (table S2). Principal component analysis (PCA) and correlation analysis indicated that the transcriptome data from the same group exhibited similar gene expression patterns and high correlation coefficients (R^2^ ≥ 0.984) (Fig. S2). Transcriptomes of *SGRi*-1 and WT plants under optimum temperature (abbreviated as CS1 for control-*SGRi*-1, and CW for control-WT), or under heat stress (HS1 for heat-*SGRi*-1, and HW for heat-WT) were compared pair-wisely (Fig. S2-S4).Venn diagram analysis showing that 972 differentially expressed genes (DEGs) were found in all four pair-wise comparisons (Fig. S3). KEGG pathway enrichment analysis showed that the enriched pathways in ‘CW vs. HW’ were significantly different from those in ‘CS1 vs. HS1’ (Fig. S4), indicating effects of heat stress and *LpSGR* knockdown were interacted at the transcriptome level. DEGs in CS1 vs. CW were mainly enriched in pathways, such as “diterpenoid biosynthesis”, “plant-pathogen interaction”, “indole alkaloid biosynthesis”, “homologous recombination”, and “zeatin biosynthesis”. It was notable that such similar KEGG pathways were also identified in HS1 vs. HW (Fig. S4).

A total of 34 DEGs involved in ROS generation and ROS scavenging pathway in *SGRi* lines and WT plants under non-stress and heat stress conditions were identified (Fig. 5I). In particular, five DEGs encoding respiratory burst oxidase homolog (Rboh, subunit of NADPH oxidase catalyzing O_2_^−^ production), including *RbohA*, *RbohB*, *RbohC*, *RbohE*, and *RbohF*, had higher transcript levels in *SGRi* lines than WT plants under heat stress. In contrast, 14 DEGs encoding peroxidase (PER), two DEGs encoding SOD, one DEG encoding CAT, three DEGs encoding APX, and three DEGs encoding GPX, and one DEG encoding DHAR had lower expression levels in *SGRi* lines than WT plants under heat stress. These DEGs clearly showed that, under heat stress, ROS-generating genes were up-regulated while ROS-scavenging enzymatic genes down-regulated in *SGRi* compared to WT (Fig. 5J).

Genes encoding light harvesting complex (LHC) and other photosystem (PS) components were greatly affected by the *LpSGR* suppression (‘CS1 vs. CW’) as well as heat stress (‘CW vs. HW’ and ‘CS1 vs. HS1’) at the transcriptional level (Fig. 6). In general, DEGs encoding photosynthetic electron transport (PET), PSⅠ, PSⅡ, LHCⅠ, and LHCⅡ proteins were down-regulated due to heat stress in both *SGRi* and WT plants, yet degrees of their expression changes were greater in *SGRi* than in WT plants (figure 6B, C). These alterations were also consistent to the immunoblot analysis of LHC proteins shown in figure 3.

**Figure 6.**
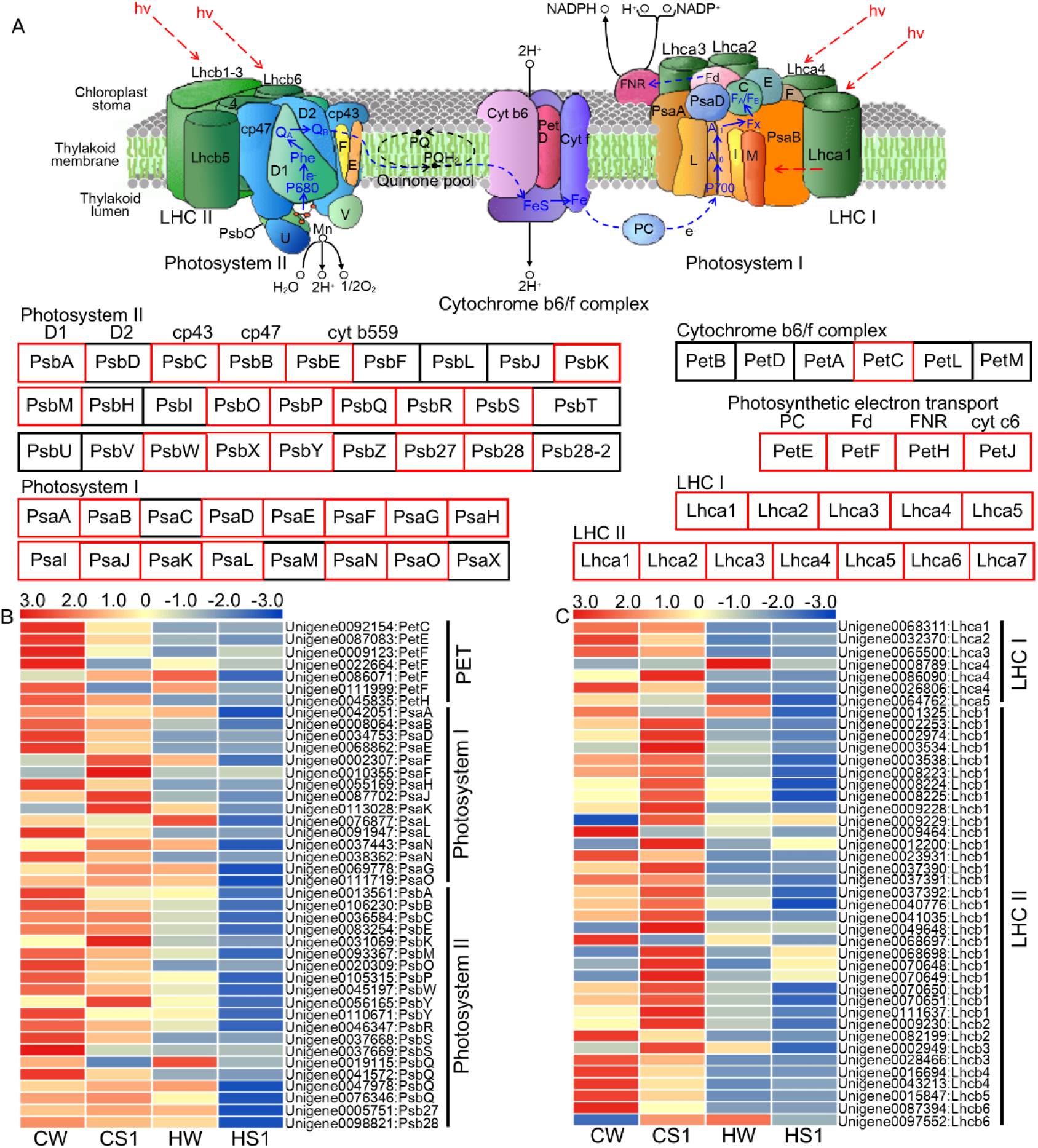
DEGs involved in photosystem and light harvesting complex. (A) Diagram of photosystem (PSⅠ and PSⅡ), light harvesting complex (LHCⅠ and LHCⅡ), and photosynthetic electron transport (PET) process, and DEGs are highlighted in red box. This diagram was drawn with reference to KEGG pathway maps (map0015 and map00196). (B) Heatmap of DEGs annotated in PET, PSⅠ and PSⅡ according to transcriptome of WT and *SGRi*-1 plants. (C) Heatmap of DEGs annotated in LHCⅠ and LHCⅡ. The RPKM of each DEGs was used for drawing heatmap. Sample abbreviations: CW, WT under control condition; CS1, *SGRi*-1 under control condition; HW, WT under heat stress condition; HS1, *SGRi*-1 under heat stress condition.

Genes coding heat stress transcription factors (HSF) was affected by knockdown of *LpSGR* as well (table S3). Under non-stress control condition, eight out of nine unigenes coding HSF, including HSFC1A, HSFA2A, HSFA2B, HSFA2C, HSFA2D, HSFB2A, and HSFB2B, except one unigen coding HSFB1, was upregulated by knockdown of *LpSGR* (table S3).

qRT-PCR analysis was performed to validate the reliability of the RNA-seq data. As shown in figure S5, the relative expression levels of 12 DEGs measured by qRT-PCR were consistent with RPKM values of transcriptomic data, supporting the validity of the transcriptome data.

### Compromised heat tolerance of *SGRi* was due to the suppression of *LpSGR* but no other long-term side effect on vegetative growth

Since constitutive suppression of *LpSGR* not only led to the cosmetic stay-green trait, reduced tiller number and therefor biomass production, but also compromised heat tolerance. It is arguable whether the compromised heat tolerance was merely a side effect due to the inhibited vegetative growth (tillering) but directly due to the suppression of *LpSGR*. Therefore, we constructed a modified Gateway-compatible ethanol-inducible binary vector by using an enhanced *alcA* promoter, with 5×alcR binding site “b”, according to Kinkema et al. (2014) and a maize ubiquitin promoter-driven *alcR* gene (Fig. S1), and generated transgenic perennial ryegrass to suppress *LpSGR* in an inducible manner (Fig. 7A). To discriminate from constitutive *SGRi* lines, we named these inducible RNAi lines as Eth-*SGRi* (figure S6).

**Figure 7.**
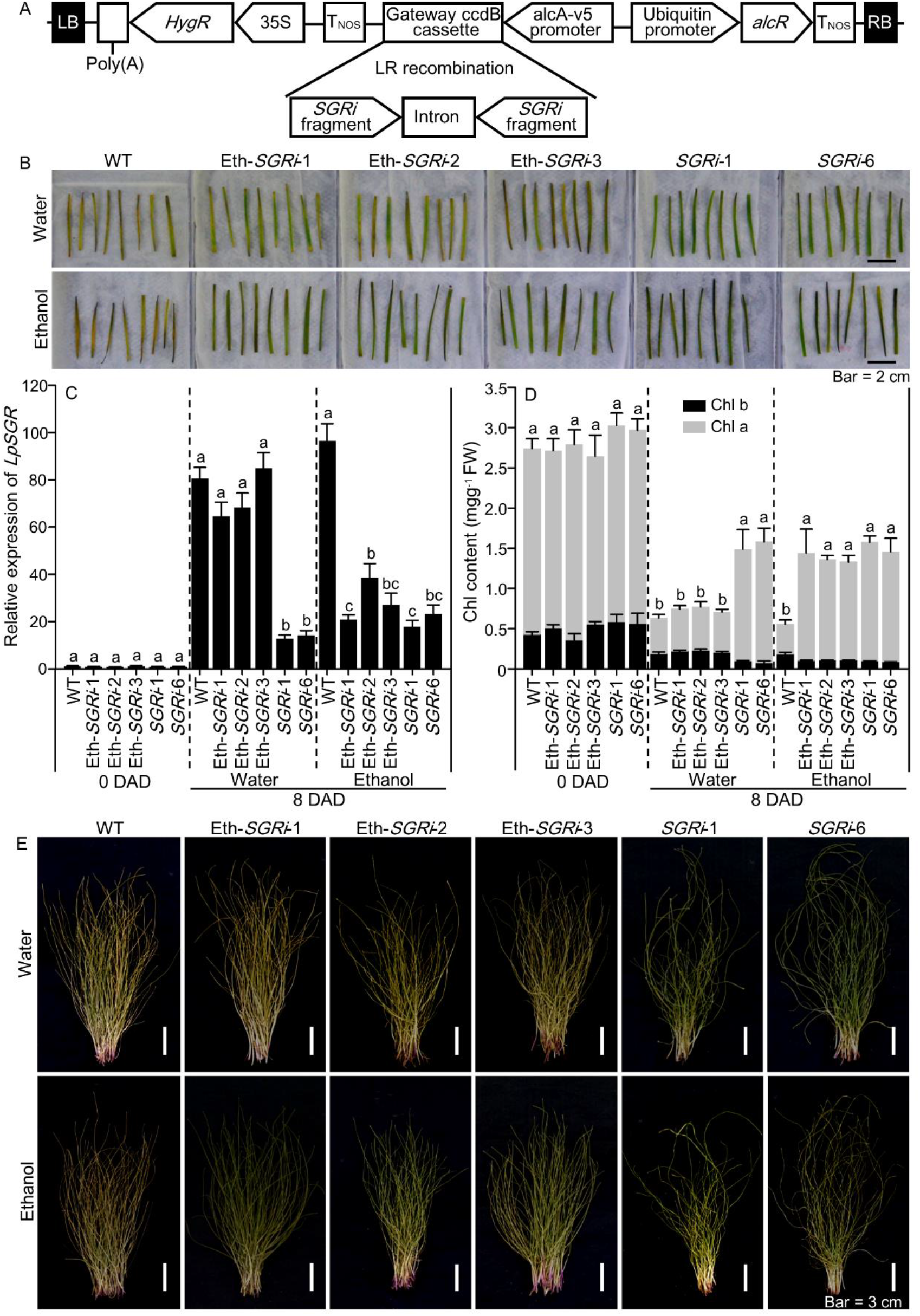
Stay-green phenotype analysis of ethanol inducible RNAi: *LpSGR* transgenic lines (Eth-*SGRi*) treated with water or ethanol under dark and post-harvest treatments. (A) Schematic diagram of ethanol inducible RNAi: *LpSGR* vector used for genetic transformation. (B-D) Phenotypes (B), relative expression levels of *LpSGR* (C) and total Chl, Chl a, and Chl b content (D) of detached leaves from WT, two *SGRi*, and three Eth-*SGRi* lines treated with water and 2% ethanol at 8 days of dark treatment. The scale bars in (B) equal to 2 cm. (E) Phenotypes of harvested above-ground biomass from WT, two *SGRi*, and three Eth-*SGRi* lines treated with water and 2% ethanol at 21 days of natural drying treatment. The scale bars in (E) equal to 3 cm. Data are means ± SE. Different letters above bars in each group represent significant difference at p ≤ 0.05.

The stringency of the ethanol inducible suppression of *LpSGR* was tested by qRT-PCR as well as phenotypic scoring. As shown in figure S7, spraying water (control) on Eth-*SGRi* lines and WT plants did not differentiate the *LpSGR* expression level in senescent leaves (4^th^ leaf from top) among these plants; while spraying 2% ethanol lead to significantly lower *LpSGR* expression levels in senescent leaves of Eth-*SGRi* lines than WT (Fig. S7). Likewise, there is no phenotypic difference between Eth-*SGRi* lines and WT plants in terms of tiller number, leaf width and length, and biomass yield (Fig. S7). Upon dark-induced leaf senescence for 8 days, *SGRi* lines showed the typical cosmetic stay-green trait, while Eth-*SGRi* lines showed the same senescence rate to WT if sprayed with water, but turned to be cosmetic stay-green if sprayed with 2% ethanol (Fig. 7B&C). We also proved that this ethanol-inducible system was effective to prevent post-harvest leaf yellowing upon ethanol treatment before harvest in the Eth-*SGRi* lines (Fig. 7D). Together, these results proved the stringency and efficacy of the ethanol-inducible system in perennial ryegrass.

Since there was no other long-term effect on vegetative growth in Eth-*SGRi* lines, we further compared their heat tolerance to WT and *SGRi* plants. As shown in figure 8, under the optimum growth temperature, there was no significant difference among these plants in terms of Chl content, Fv/Fm, Pn, and EL of leaves treated with water or 2% ethanol. Under heat stress, when all plants were sprayed with water, Eth-*SGRi* lines and WT had similar values of the above-mentioned physiological parameters which were all significantly higher than those of *SGRi* lines; when sprayed with ethanol, Eth-*SGRi* lines also showed compromised heat tolerance with significantly lowers values of these physiological parameters than WT (Fig. 8). Together, these results showed that the compromised heat tolerance trait of *SGRi* and Eth-*SGRi* lines was solely dependent on the suppression of *LpSGR* but no other long-term effect on vegetative growth.

**Figure 8.**
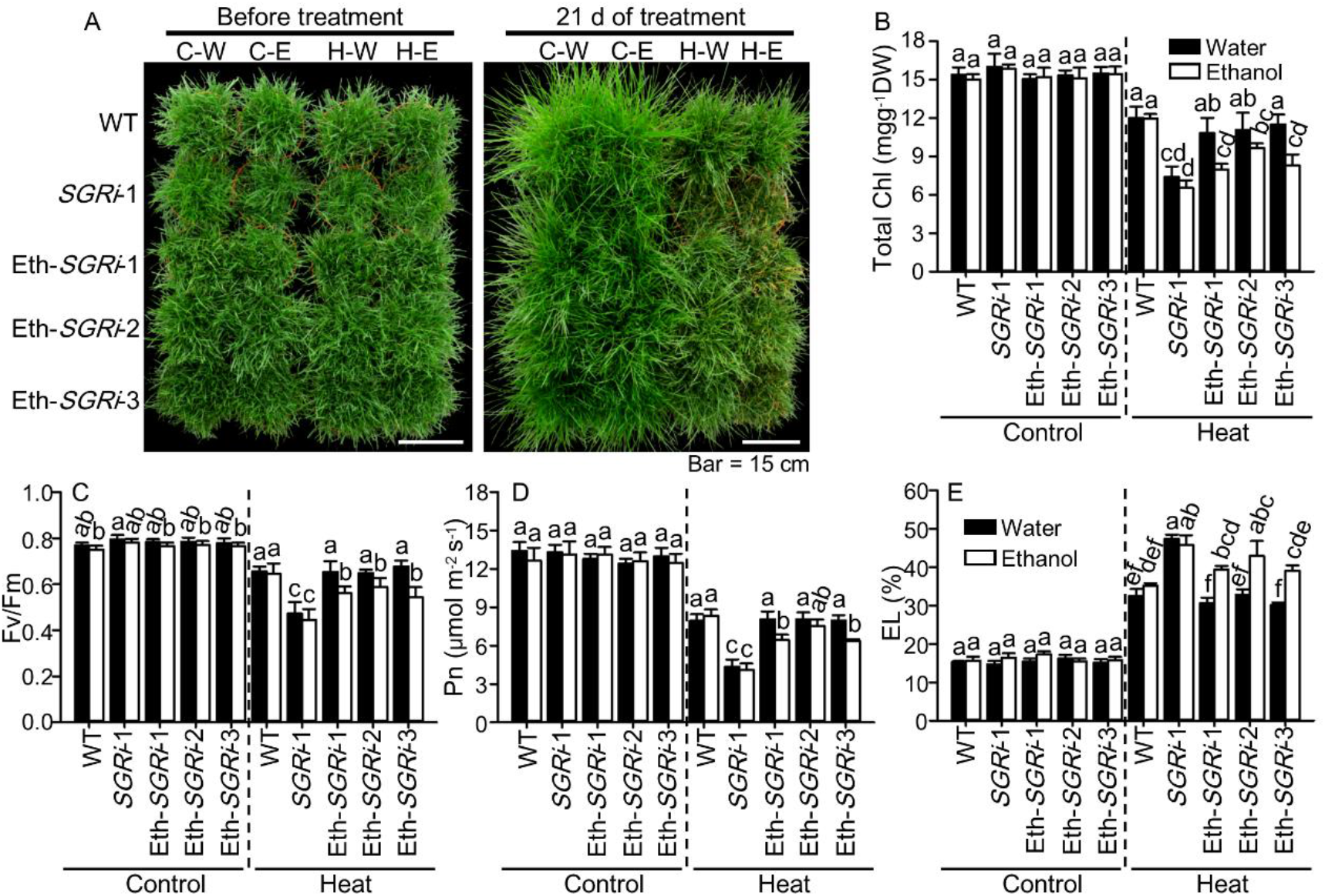
Phenotypic and physiological analysis of Eth-*SGRi* transgenic lines treated with water or ethanol under heat and non-stress conditions. (A-E) Phenotypes (A), total Chl content (B), Fv/Fm (C), Pn (D), and EL (E) of WT, *SGRi*-1, and Eth-*SGRi* lines treated with water and 2% ethanol at 21 days of heat or non-stress treatment. The scale bars in (A) equal to 15 cm. Data are means ± SE. Different letters above bars in each group represent significant difference at p ≤ 0.05.

## Discussion

Chl molecules play critical roles in absorbing light energy for photosynthesis, which is essential for plant growth and development (Tanaka & Tanaka, 2006; Morita et al., 2009). Disrupting Chl a degradation by mutating or knockdown *SGR* led to stay-green phenotype during dark-induced leaf senescence in many plant species (Armstead et al., 2007; Jiang et al., 2007; Park et al., 2007; Ren et al., 2007; Zhou et al., 2011; Shimoda et al., 2016). However, occasional accumulation of free Chl molecules or Chl catabolic intermediates can cause photodamage to chloroplast in plant cells or phototoxicity to seed (Hirashima et al., 2009; Li et al., 2017). In this study, using both constitutive or ethanol inducible suppression of *LpSGR*, we provided evidence that knockdown of *SGR* exacerbated heat stress-induced photodamage and further accelerated stability loss of chloroplast and cell death rather than maintained the stay-green phenotype in perennial ryegrass.

### *LpSGR* knockdown transgenic lines had no stay-green phenotype under heat stress

The stay-green trait is defined by the retention of Chl during senescence (Hörtensteiner, 2013). Stay-green mutants are further divided into two types, that is functional stay-green and cosmetic stay-green mutants. The functional stay-green mutants show delayed initiation of senescence or slower progression of senescence than their reference plants, while the cosmetic stay-green mutants show normal progression of senescence but remains greenness during leaf senescence. Null mutants in *sgr* were responsible for Mendel’s green pea (Armstead et al., 2007), and were identified in rice (*Oryza sativa*) (Jiang et al., 2007; Park et al., 2007), and *Arabidopsis thaliana* (Ren et al., 2007), all of which showed cosmetic stay-green.

Besides the stay-green phenotype, long term effect of Chl retention on plant phenotype in *sgr* mutant also include ineffective nitrogen remobilization (Macduff et al., 2002). Knockdown of *SGR* in perennial ryegrass caused increased protein content in the aboveground biomass (Xu et al., 2019), and mutation of *SGR* in the leguminous plant *Medicago truncatula* also affected root nodule formation and senescence rate (Zhou et al., 2011). The role of *SGR* in plant nitrogen remobilization during senescence or sink/source conversion process should contribute to the phenotypic alternation observed in constitutive suppression of *SGR* in perennial ryegrass for the reduced tiller numbers and aboveground biomass accumulation.

In addition to the side effect on nitrogen remobilization in the *sgr* mutant, double mutation of *SGR1* and *SGR2* caused a severe phototoxic injury to maturing seeds in *Arabidopsis thaliana* and thus reduced seed longevity (Li et al., 2017). In this study, we found that *SGRi* lines not only lost the stay-green phenotype under heat stress but were more heat susceptible than WT. The rationale behind this observation was likely due to disrupted electron chain transport, quantum yield with energy dissipation, ROS yield in chloroplast and ROS-scavenging systems when under heat stress discussed as follows.

### Involvement of *SGR* in chloroplast development including grana stacks’ structure and photosynthetic proteins stability

SGR is an essential Chl catabolic enzyme (CCE) for it interacts directly or indirectly with all major CCEs and may play a role in recruiting CCEs onto thylakoid membranes (Sakuraba et al., 2012). In the current study, ultrastructural analyses of *SGRi* and WT plants showed several notable differences in grana stacks’ structure under both non-stress control and heat stress conditions. Firstly, the grana number *per* chloroplast were significantly higher and granal lamellaes were more tightly stacked in *SGRi* lines than those in WT under non-stress condition, which could contribute to the significantly higher quantum yield of PSⅡ and apparent photosynthetic electron transport rate in *SGRi* lines than that in WT under non-stress condition. Secondly, the grana size was smaller in *SGRi* lines than in WT under non-stress condition. One recent report revealed that *SGR* is involved in the formation of PSⅡ (Chen et al., 2019), that explains the altered chloroplast grana development in *SGRi* plants. This altered grana stacks’ structure in *SGRi* was not reported in other mutants of Chl catabolic genes yet (e.g. *nyc1*, *pph*, and *nyc3* mutants) (Kusaba et al., 2007; Morita et al., 2009; Schelbert et al., 2009). Thirdly, the thylakoid membranes and LHC proteins were more greatly degraded in all *SGRi* lines than in WT under heat stress, which was similar to the results of dark-induced rapid cell death in *Accelerated cell death1* (*Acd1*, also known as *pao*) mutant (Hirashima et al., 2009), and to the light-induced photodamage to Arabidopsis seeds of *sgr1*/*sgr2* double mutant (Li et al., 2017). Together with previous findings, our results suggest that rapid turnover of Chl a, that is primarily catabolized by SGR and possibly affected by the further downstream catabolic enzyme PAO (Hirashima et al., 2009), was important to minimize PS system damage under heat or other stress condition (e.g., photodamage to seed).

### Involvement of *SGR* in electron transport and energy dissipation

The light energy absorbed by PSⅡ antennae can be either utilized via photochemistry or dissipated via various thermal processes in a competitive manner. High or low temperature can induce changes in energy partitioning in the photosynthetic apparatus (Hendrickson et al., 2004). Partition of different routes of the excitation energy utilization/dissipation in PSⅡ complexes (energy partitioning) is an important response of photosynthetic apparatus to environmental factors (Agrawal & Jajoo, 2021; Kalaji et al., 2016; Wang et al., 2009). Chl fluorescence parameters, including Y(Ⅱ), Y(NPQ), Y(NO), and ETR, provide a rapid, efficient and non-invasive means to investigate photosynthetic processes and to detect environmental effects on photosynthetic apparatus (Agrawal and Jajoo, 2021; Wang et al., 2009). As for the quantum yield parameters, Y(Ⅱ) represents the effective PSⅡ quantum yield, Y(NPQ) is the quantum yield for dissipation by downregulation as a protective mechanism against excessive light energy, Y(NO) represents the quantum yield for non-regulated energy dissipation, and the sum of Y(Ⅱ), Y(NPQ), and Y(NO) equals to one. In the current study, the Y(Ⅱ) and ETR values were significantly higher, but significantly lower in *SGRi* lines than those of WT plants under non-stress and heat stress conditions, respectively. These results indicated that, under non-stress condition, the absorbed quanta are more converted into chemically fixed energy by the photochemical charge separation and high rate of charge separation at PSⅡ reaction centers in *SGRi* lines, but *vice versa* when under heat stress. The Y(NPQ) value in *SGRi* lines was significantly lower than that of WT plants under non-stress, but higher than WT when under heat stress. The higher value of Y(NPQ) indicates that leaves are receiving excessive the photon flux density and reflects that the plant with high Y(NPQ) protects itself by regulating the dissipation of excessive excitation energy into harmless heat (Wang et al., 2009). In contrast to Y(NPQ), the Y(NO) value indicates excessive and unregulated energy dissipation, e.g., excessive energy causing ROS generation (Wang et al., 2009). *SGRi* lines had similar level of Y(NO) to WT at optimal temperature, but higher level of Y(NO) than WT under heat, suggesting that shutting down SGR and Chl a catabolism caused inefficient photochemical energy conversion and non-regulated energy dissipation, which in turn favors the production of ROS under heat stress.

### Involvement of *SGR* in ROS balancing under heat stress

In general, heat stress can disrupt the balance between light energy absorption and utilization for photochemical electron transport causing photoinhibition and photodamage of PSⅡ (Rossi et al., 2021; Wang et al., 2009). And shutting down SGR and Chl a catabolism further exacerbated the degree of non-regulated energy dissipation from the PS system (Fig. 4). In photosynthetic organism, energy absorbed by PSⅡ is normally consumed for driving photosynthetic electron transport for ATP and NADPH synthesis. When the excitation energy transferred to PSⅡ exceeds that can be utilized, the excessive energy can be dissipated through non-photochemical quenching (NPQ) or transferred to O_2_ to produce ROS (Wilson et al., 2006). And the burst or accumulation of ROS further causes photodamage onto PSⅡ reaction centers (Wilson et al., 2006). Under non-stress condition, *SGRi* lines had higher quantum yield of PSⅡ [Y(Ⅱ)] as well as higher ETR than WT possibly due to altered Chl a/b ratio and chloroplast structure. Yet, this high efficiency of ETR and Y(II) could lead to higher susceptibility to heat stress in *SGRi* lines. Thus, under heat stress, the majority of absorbed energy was converted into non-regulated energy, which explains the higher concentrations of ROS (O_2_^−^ and H_2_O_2_) in *SGRi* lines.

The production of O_2_^−^ can be catalyzed by respiratory burst oxidase homologue (Rboh, a subunit of NADPH oxidase) proteins, and O_2_^−^ can be convert to H_2_O_2_ dismutated by SOD (Liu et al., 2016; Waszczak et al., 2018). In *Arabidopsis*, ten genes encoding Rboh proteins have been identified (Liu et al., 2016). In this study, five DEGs encoding Rboh, including RbohA, RbohB, RbohC, RbohE, and RbohF, had higher transcript levels in *SGRi* lines than WT under heat stress, that is consistent to the elevated ROS levels in *SGRi* lines.

The accumulated ROS can cause damage to proteins, lipids, DNA, and carbohydrates and ultimately lead to senescence and cell death (Apel & Hirt, 2004; Waszczak et al., 2018). The ROS scavenging system in higher plants contains both enzymatic and non-enzymatic components (Zhang et al., 2016a). The activation of enzymatic antioxidant systems or accumulation of non-enzymatic compounds can suppress over-production of ROS and benefit plant stress tolerance (Apel & Hirt, 2004; Sachdev et al., 2021). Under heat stress, activities of SOD, CAT, APX, and GPX in *SGRi* lines were significantly lower than those in WT plants under heat stress, which is consistent to down-regulated expression of their encoding genes according to the transcriptome analysis.

Based on these results, we proposed that blocking *SGR* leads to more efficient quantum yield and ETR, and smaller but more stacked grana/thylakoid at optimum temperature, but heat stress induced excess energy dissipation in both regulated and non-regulated manners. The excessive non-regulated energy dissipation further resulted in burst and accumulation of ROS, causing severe degradation of LHC proteins and the thylakoid system in *SGRi* plants.

### Application potential of ethanol inducible system in forage production

Constitutive overexpression or suppression of genes has been the traditional approach for characterize gene function or genetic improvement in plants. However, this approach is somehow limited when the target genes are detrimental to plant development or have multiple side effects (Laufs et al., 2003). Chemical inducible gene expression system can be used to analyze gene function more precisely, by inducing target gene(s)’ expression with insecticides, steroids, estrogens, antibiotic tetracycline, and ethanol (Laufs et al., 2003; Moore et al., 2006). Ethanol is affordable, biodegradable, readily available, environmentally friendly, and non-phytotoxic at the levels necessary to induce gene expression (Kinkema et al., 2014). Therefore, ethanol inducible gene expression systems hold great promise for genetic improvement in forages and other cash crops.

The stay-green phenotype of *SGRi* lines is a highly desirable trait for perennial grass used as turf and forage. However, constitutive suppression of *LpSGR* decreased its biomass production and heat tolerance. Based on the enhanced *alcA* promoter according to Kinkema et al. (2014), we further modified it into a Gateway-compatible and ethanol-inducible binary vector by using a maize ubiquitin promoter to drive the *alcR* gene and a ccdB cassette driven under the modified alcA-V5 (version 5) promoter to facilitate gene cloning using the one step LR recombination. Our results showed that the Eth-*SGRi* lines had no phenotypic difference from the WT plants in terms of tiller number, leaf width and length, biomass yield, and heat tolerance if only not induced by ethanol. Prior to harvest, once the Eth-*SGRi* lines were sprayed with ethanol, these lines showed the same stay-green trait to *SGRi* lines that is highly desirable for forage purpose. Thus, these results not only proved the stringency and efficacy of the ethanol-inducible system in perennial ryegrass, but also holds a promise for its application in forage grass breeding in the future.

### Conclusion

In summary, Chl a catabolism is essential for plant heat tolerance that blocking this catabolic step by suppressing *SGR* caused severe heat-induced phototoxic injury with disrupted photosystem and activated ROS generation in perennial ryegrass. As illustrated in figure 9, under optimum temperature, suppressing *SGR* leads to the typical stay-green phenotype with Chl retention in senescent leaves; while under heat stress, suppressing *SGR* leads to compromised heat tolerance with more rapid leaf senescence than WT plants. This compromised heat tolerance in *SGRi* plants can be remedied by using ethanol inducible system to avoid side effect of *SGR* suppression but to keep the stay-green trait in forage production.

**Figure 9.**
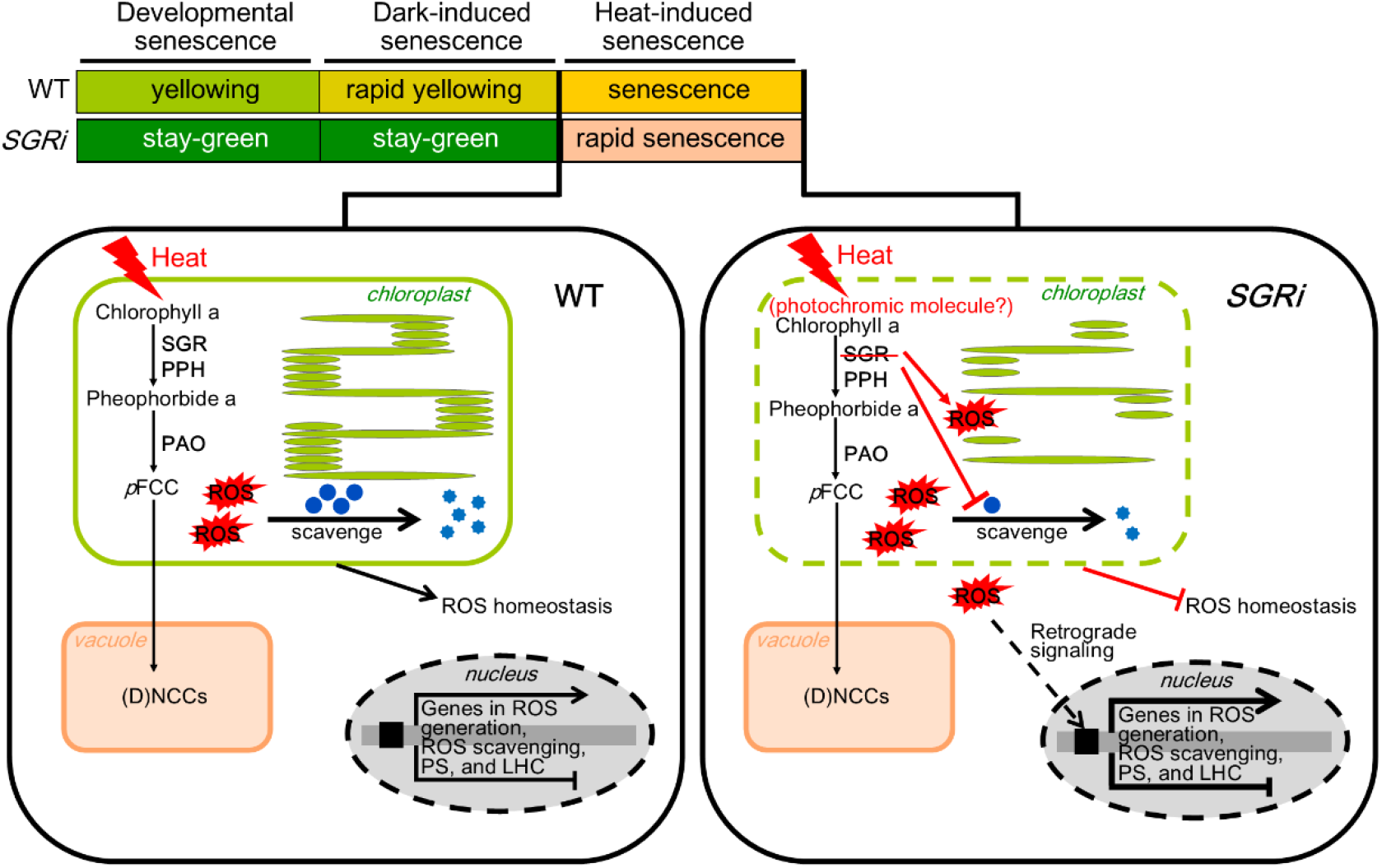
Proposed model illustrating the influence of blocking *SGR* on plant heat tolerance. Under developmentally senescence and dark-induced senescence condition, the *SGRi* leaves show a stay-green phenotype, while under heat stress condition, the *SGRi* leaves show an accelerated or rapid senescence phenotype. This model was drawn with reference to Aubry et al. (2020), Liu et al. (2021), and Sakuraba et al. (2014).

## MATERIALS AND METHODS

### Construction of Gateway-compatible and ethanol inducible RNAi vector

The ethanol inducible gene expression system consists of the AlcR transcription factor and its target promoter, *alcA* (Felenbok, 1991). AlcR binds to specific site in the *alcA* promoter and, in the presence of ethanol, initiates downstream gene’s transcription. The modified *alcA_V5* promoter with higher specificity and activity (Kinkema et al., 2014) was used to drive the Gateway cassette of attR1-*ccdB*-attR2-*T*_*NOS*_. The maize ubiquitin promoter-*AlcR-T*_*NOS*_ cassette and *alcA_V5-ccdB* cassette were synthesized by a gene synthesis company (GENEWIZ Inc. Suzhou, China), and then insert into the multiple cloning site (MCS) of pCAMBIA1300 to generate the new Gateway-compatible and ethanol inducible vector pCAMBIA1300-UbiP-*alcR*-*alcA_V5*-*ccdB* (Fig. S1), through which the target gene can be readily subcloned into the vector under driven of the *alcA_V5* promoter.

The same fragment of *LpSGR* (Xu et al. 2019) was used to generate the RNAi hairpin structure. In short, the entry vector pGMKannibal-2×*LpSGR* and the destination vector of pCAMBIA1300-UbiP-*alcR*-*alcA_V5*-*ccdB* was recombined through LR reaction (Invitrogen, Carlsbad, CA, USA) to generate pCAMBIA1300-UbiP-*alcR*-*alcA_V5*-2×*SGRi* (Fig. S1). The generated vector was verified by vector size and enzyme digestion for the presence of 2×*SGRi* fragment and then electro-transformed into *Agrobacterium tumefaciens* strain ‘AGL1’ cells.

### Genetic transformation of perennial ryegrass

Perennial ryegrass cv ‘Buena vista’ was used to generate transgenic lines. *Agrobacterium tumefaciens*-mediated transformation was carried out to generate stable transgenic perennial ryegrass, according to the protocol reported before (Xu et al., 2019; Zhang et al., 2013). Putative transgenic plants were confirmed by PCR verification for the presence of the *HPTII* gene and qRT-PCR analysis of target gene’s relative expression level. The ethanol-inducible stable transgenic lines were named as Eth-*SGRi* lines, in contrast to the constitutive RNAi lines driven under the maize *ubiquitin* promoter, named as *SGRi*, as reported before (Xu et al., 2019).

### Plant materials, growth conditions and treatment

The non-transgenic (wildtype, WT) and transgenic lines (*SGRi* lines and Eth-*SGRi* lines) of the same variety ‘Buena vista’ were compared in this study. All plants were grown in a mixture of peat, vermiculite, and perlite (3:3:1 v/v) in plastic pots at photoperiod of 14/10 hr, 25/20°C, day/night except for heat stress, relative humidity of 70%, and light intensity of 750 μmol photos m^−2^ s^−1^. Plants were watered and fertilized weekly with half-strength Hoagland’s nutrient solution (Hoagland & Arnon, 1950).

For testing whether knockdown of *LpSGR* may affect plant phenotypic traits, the single tillers of *SGRi*, Eth-*SGRi* and WT and were split from the mother plants and grown under the same condition. After one month of growth, the plant height, width and length of mature leaf (3^rd^ leaf from the top), tiller number, and shoot fresh weight of each plant were measured.

For heat stress treatment, ryegrass plants were vegetative propagated and maintained at a height of 12 cm by weekly mowing. After two-months growth in the growth chamber, plants were either grown under the optimum temperature at 25/20°C (day/night) or exposed to high temperature at 38/35°C (day/night) in growth chambers for 21 days. The photoperiod, relative humidity, and light intensity were set at the same values as described above. As for the ethanol treatment, ryegrass plants were treated with water (control) or 2% ethanol one week prior to and once a week during the heat stress treatment.

For dark-induced leaf senescence, the excised mature leaves (3^rd^ leaf from the top) were wrapped in paper towels moistened with 3 mM 2-(N-morpholino) ethanesulfonic (MES) buffer (pH 5.8), with or without 2% (v/v) ethanol, in the dark at 25°C for 8 days (Zhang et al., 2016b). For the forage post-harvest storage test, the Eth-*SGRi*, *SGRi*, and WT plants were pretreated with 2% ethanol or water control for one week prior to harvest, then the harvested shoot were dried for 21 days under natural condition.

### Measurement of physiological parameters

Chl content was determined following dimethyl sulfoxide (DMSO) extraction protocol described by Barnes et al. (1992). Membrane stability was evaluated by measuring electrolyte leakage (EL) following a method described by Blum and Ebercon (1981). In brief, 0.1 g of leaves were immersed in 30 ml distilled deionized water in a 50 ml centrifuge tube with constant shaking for 24 hr at room temperature. The initial conductance reading (Ci) of the incubated solution was taken using a conductivity meter (Thermo Scientific). Then the leaf samples were autoclaved at 121 °C for 20 min and shaking for another 24 hr, and then the maximum conductance (Cmax) of the solution was measured. The relative EL was calculated as Ci/Cmax × 100. Net photosynthesis rate (Pn) was measured using Li-COR6400 portable photosynthesis system (LI-COR Inc., Lincoln, NE, USA) according to the protocol described by Burgess and Huang (2014). The leaf relative water content (RWC) was measured by previously described protocol (Zhang et al., 2020).

The Chl fluorescence parameters were measured with Chl fluorometer PAM-2500 (Heinz Walz GmbH, Effeltrich, Germany). In brief, leaves were dark adapted for 15 min prior to measurement. The minimum fluorescence (Fo) was measured by applying 1 Hz light pulses. The maximum fluorescence (Fm) was obtained by applying 10 Hz saturating blue pulse. The maximum fluorescence yield (Fm’) and the Chl fluorescence yield (Fs) was determined by switching on 183 μmol m^−2^ s^−1^ actinic irradiance and saturating pulses were then applied at 30 s intervals for 5 min. The values of various Chl fluorescence parameters were captured at each saturation pulse interval. Photosynthetically active radiation (PAR) on the leaf was applied at 0-3702 μmol m^−2^ s^−1^ in rapid light curve. Maximum photochemical efficiency of PSII (Fv/Fm), quantum yield of PSⅡ [Y(Ⅱ)], quantum yield of regulated energy dissipation [Y(NPQ)], quantum yield of non-regulated energy dissipation [Y(NO)], photochemical fluorescence quenching (qP), and apparent photosynthetic electron transport rate (ETR) were calculated based on the key parameters mentioned above using the equations as previously described by Wang et al. (2009).

The methods for quantification of O_2_^−^ and H_2_O_2_ production, histochemical staining for O_2_^−^ and H_2_O_2_, and activity analysis of antioxidant enzyme, including SOD, CAT, APX, and GPX, were previously described by Zhang et al. (2016a).

### Transmission electron microscopy

Transmission electron microscopy was performed according to the methods described by Morita et al. (2009) with slight modifications. The middle section (~1 cm) of mature leaves were excited and firstly fixed with 2.5% glutaraldehyde in 100 mM cacodylate buffer (pH 7.4). The Samples were post-fixed with 1.5% OsO_4_ in 100 mM cacodylate buffer (pH 7.4) for 90 min. Leaf samples were then embedded in Quetol resin mixture (Quetol 812, Nisshin EM) and sectioned using an ultramicrotome. The sections were mounted on copper grids and stained with 3% uranyl acetate and lead citrate. Micrographs of chloroplast were observed using a transmission electron microscope (Hitachi, H-7650).

### Immunoblotting

Protein was extracted by homogenization of 100 mg leaf samples in 1.0 ml protein extraction buffer (50 mM Tris-HCl buffer containing 2.5% [w/v] SDS, 10% [w/v] glycerin, 2.5% [v/v] β-Mercaptoethanol, pH 6.8). After 20 min incubated on ice, the homogenate was centrifuged at 12000 rpm for 10 min at 4 °C and the supernatant was saved for protein content and immunoblotting analyses. The protein content in the supernatant was determined as described by Bradford (1976). Ten μl of protein solution was subjected to SDS-PAGE (12% polyacrylamide gel, made by Invitrogen) and the electrophoresed proteins were then transferred onto a polyvinylidene difluoride membrane (Immobilon^®^-P^SQ^ membrane, Millipore GmbH, Eschborn, Germany) for immunoblotting analysis. The target proteins were detected with primary antibodies, including Lhca3, Lhcb1, Lhcb2, Lhcb3, Lhcb5, PsaA, PsbA (D1), PsbD (D2), and RbcL, from Agrisera (http://www.agrisera.com/) and secondary HRP-conjugated goat anti-rabbit IgG antibody form Invitrogen (Shanghai, China). Target proteins in PVDF membrane were reacted with SuperSignal™ West Femto Maximum sensitivity substrate (Thermo Scientific) and then were visualized using the Fusion Solo chemiluminescence system (VILBER LOURMAT, France).

### Transcriptomic analysis

Leaves of *SGRi*-1 and WT plants grown under the optimum temperature or 21 days of high temperature and treatment were harvested and quickly frozen in liquid nitrogen. Leaves from each pot of plants were regarded as one biological replicate, and three biological replicates were performed for RNA-seq analysis in each treatment. Total RNA isolation, cDNA library construction, Illumina sequencing, data processing, and differentially expressed gene (DEGs) analysis (i.e. GO and KEGG analysis) were carried out by a commercial gene sequencing company (Gene Denovo Corporation, Guangzhou, China) and were the same as described in our previous study (Xu et al., 2019). The sequencing data are available from the NCBI bioproject (accession No. PRJNA756535).

### qRT-PCR analysis

The total RNA was isolated using Plant RNA Kit (Omega Bio-tek, Georgia, USA). After DNA digestion with Perfect Real Time gDNA Eraser Kit (TaKaRa, Otsu, Japan), first strand cDNA was synthesized using PrimeScript™ RT reagent Kit (TaKaRa). The qRT-PCR reaction was performed in a 20 μl reaction volume using SYBR green master mix (Thermo Scientific) on a Roche LightCycler^®^ 480 II Real-Time PCR machine. The qRT-PCR analysis was performed with four biological replicates and relative expression levels of target genes were calculated using the 2^−ΔΔCT^ method with *LpeIHF4A* as the reference gene (Huang et al., 2014). Detailed information of primers used in this study are listed in Table S1.

### Statistical analysis

Data from all samples were statistically analyzed using SPSS software (Version 12, SPSS Inc., Chicago, IL) and verified by Duncan test at a significance level of 0.05. The data are expressed as means ± SE.

## SUPPLEMENTAL DATA

**Supplemental Figure S1. Construction of ethanol inducible RNAi vectors.** To construct the Gateway-compatible and ethanol-inducible binary vector, pCAMBIA1300-UbiP-alcR-alcA-ccdB, we modified pCAMBIA1300 by inserting the ubiquitin promoter-*AlcR-T*_*NOS*_ cassette and alcA promoter-attR1-*ccdB*-attR2-*T*_*NOS*_ cassette into the multiple cloning site (MCS). The Eth-SGRi vector was constructed by LR reaction by recombining the pEnD-Kannibal-2xSGRi vector (Xu et al., 2019) with the binary vector, to generate pCAMBIA1300-UbiP-*alcR*-alcA-2x*SGRi*.

**Supplemental Figure S2. Principal component analysis (PCA) of gene expression patterns and correlation analysis of genes from the same group.** (A) Principal component analysis (PCA) of gene expression patterns. PC1, the first principal component; PC2, the second principal component. (B) Correlation analysis of genes from the same group.

**Supplemental Figure S3. Count and Venn diagram of DEGs according to transcriptomes of WT and *SGRi*-1 plants under non-stress control and heat stress conditions.** (A) Count of DEGs among each group. (B) Venn diagram of DEGs in the compared groups. Numbers in each overlapping part represent DEGs that were co-expressed in multiple stages.

**Supplemental Figure S4. KEGG pathway enrichment of DEGs in the pair-wise comparison of groups.** (A-D) KEGG pathway enrichment of DEGs in the pair-wise comparison of transcriptomes of CW vs. CS1 (A), CW vs. HW (B), CS1 vs. HS1 (C), and HW vs. HS1 (D).

**Supplemental Figure S5. RNA-seq validation using the qRT-PCR analysis.** The black bars represent RAN-seq data and were built using RPKM of selected DEGs; the white bars represent qRT-PCR results. Data are means ± SE.

**Supplemental Figure S6. Characterization of Eth-*SGRi* lines.** (A) PCR confirmation of transgenic lines (lines: Eth-*SGRi*-1, −2, and −3) by amplifying a fragment of *HPTII* gene. (B) qRT-PCR analysis of *LpSGR* in senescent leaves (4^th^ leaves from top) of Eth-*SGRi* lines and WT when treated with water and 2% ethanol. Data are means ± SE. Different letters above bars in each group represent significant difference at p ≤ 0.05.

**Supplemental Figure S7. Growth traits of WT, *SGRi* lines and Eth-*SGRi* lines after 2-months of growth under non-stress condition.** (A) Growth phenotypes of WT, *SGRi* and Eth-*SGRi* lines before and after 2-months of growth under non-stress condition. The scale bars equal to 7.5 cm. (B-E) Mature leaf length (B), mature leaf width (C), tiller number (D), and shoot fresh weight (E) of WT, *SGRi*-1, and Eth-*SGRi* lines. Data are means ± SE. Different letters above bars in each group represent significant difference at p ≤ 0.05.

**Supplemental Table S1. Primers used in this study.**

**Supplemental Table S2. The assembled results and gene annotation in transcriptome in WT and SGRi plants.**

**Supplemental Table S3. Differentially expressed *HSF* genes identified in the pair-wise comparison of groups.** ‘-’represent target genes which were not differentially expressed in specific group.

